# Forensic investigative genetic genealogy match rate estimated from a nation-wide population register

**DOI:** 10.64898/2026.08.06.743282

**Authors:** Harel Ismach, Gili Greenbaum, Debbie Kennett, Shai Carmi

**Affiliations:** Braun School of Public Health and Community Medicine, The Hebrew University of Jerusalem, Jerusalem, Israel; Department of Ecology, Evolution and Behavior, The Hebrew University of Jerusalem, Jerusalem, Israel; Research Department of Genetics, Evolution and Environment, University College London, London, United Kingdom

## Abstract

Forensic investigative genetic genealogy (FIGG) is a revolutionary method in forensic genetics, whereby genetic relatives of an unknown target person are detected in direct-to-consumer genomic databases; the family trees of these relatives are then reconstructed to suggest candidates for the target person. Despite recent successes, the full potential of the technology has not yet been systematically evaluated at the scale of an entire country. To estimate the proportion of FIGG cases where a genetic relative is detected in the database (the “match rate”), we used the Israeli population register, covering all past and present citizens. After extensive quality control, the register included 12.4 million individuals, among them 10.4 million alive. We simulated genomic databases by randomly selecting subsets of predefined sizes of the live adult population. With a database covering 1% of the population, we estimate that FIGG would detect at least one relative of fifth degree (e.g., a second cousin) or closer for about 25% of the population, and at least two relatives for 9% of the population. A database covering 5% of the population would find at least one relative of third degree (e.g., a first cousin) or closer for half the population. More distant relatives are rarely identified in the register, likely due to its limited time depth. Our results provide the first country-wide direct estimates for the utility of FIGG in generating investigative leads.

## Background

Forensic investigative genetic genealogy (FIGG), available to law enforcement since 2018, is one of the most powerful tools ever developed for forensic identification. FIGG became possible with the rapid expansion of direct-to-consumer genomic databases, which now include the genome-wide profiles of tens of millions of individuals. To identify a target person based on DNA extracted from a forensic sample, investigators generate a genome-wide profile for the target (using whole-genome sequencing or microarray genotyping), upload the profile to one of the databases where upload is permitted, and search for relatives (“matches”). Genealogists then reconstruct the family trees of each of the matches, attempting to intersect the trees and identify the target person^1–6^.

FIGG has had an enormous impact since its inception. Based on one database, as of June 2026, over 1300 cases have been solved^7^. One commercial lab reported closing more than 7,600 years of investigation, including 121 identifications in cases >30 years old^8^, and another reported solving over 400 cases^9,10^. However, the general success rate of FIGG, or the proportion of cases solved, has not been reported by forensic labs. Nevertheless, such estimates, even for specific components of FIGG, are crucial for developing evidence-based policy, particularly given the ethical, social, and legal complexities of FIGG^5,11^.

Previous studies used multiple approaches to estimate FIGG’s success rate. Population genetic modeling^2,12,13^ and large-scale genomic data^2^ predicted that FIGG would succeed in finding a match to an unknown target for about 40-60% of the targets given realistic database sizes. One commercial lab reported detecting a third cousin or a closer match in 80% of their cases^4^. However, these results do not account for difficulties with the genealogical search *after* a match has been found. Particularly, matches might not always identify under their real names, public records often do not cover all of their ancestors, genealogical databases might be subject to variable access policies^14,15^, and an intersection between the family trees of matches might be difficult or impossible to find. Thus, the target person may not always be reachable from the match based on public genealogical data. Empirical identification attempts^16–18^ suggested success rates of around 40-70%, but published studies were small. In addition, the empirical studies used existing international databases, not addressing the potential of dedicated national genomic initiatives.

Here, we leverage a national population register to provide realistic estimates of FIGG’s match rate, namely the proportion of unknown target individuals who have a close relative in a genomic database. Our modeling approach, based on a national register, allows us to focus on *genealogically documented* relatives. Specifically, to simulate FIGG, we assign a random subset of the population to the genomic database, and we then record the fraction of the population that has one or more relatives in the database. Importantly, in our approach, relatedness is inferred from the genealogy implied by the population register, without relying on public or commercial genealogical records.

## Results

To estimate the FIGG match rate, we used deidentified data from Israel’s Population and Immigration Authority (IPIA). Each IPIA record includes the ID of one citizen, whether or not deceased, sex, year of birth, and the IDs of the two parents (see Methods). The IPIA data covered 12,563,620 individuals, among them 10,498,709 alive as of 2021. After extensive quality control (see Methods), 105,895 individuals were excluded from downstream analyses. Basic statistics of the IPIA data are shown in Table 1, Table S1, Figure 1, and Figure S1.

**Table 1.** Statistics of the IPIA data. The dataset initially included 12,563,620 individuals. The statistics shown are after exclusion of 105,895 individuals who failed quality control (see Table S1 for all statistics before and after quality control). SD: standard deviation.

|  |  |
| --- | --- |
| Total number of individuals | 12,457,725 |
| Number of individuals alive | 10,430,877 |
| Number of deceased individuals | 2,026,848 |
| Number of males | 6,262,793 |
| Number of females | 6,194,932 |
| Number of individuals with each number of parents recorded | 0 parents: 3,465,674<br>1 parent: 689,477<br>2 parents: 8,302,574 |
| Number of individuals with one or more half-siblings | 685,856 |
| Fraction of individuals without children | 55.24% |
| Mean (SD) of the number of children per individual | All individuals: 1.38 (2.04)<br>Only individuals with children: 3.09 (2.01) |
| Mean (SD) of the paternal age at birth | 32.5 (6.78) |
| Mean (SD) of the maternal age at birth | 28.7 (5.82) |

**Figure 1.**
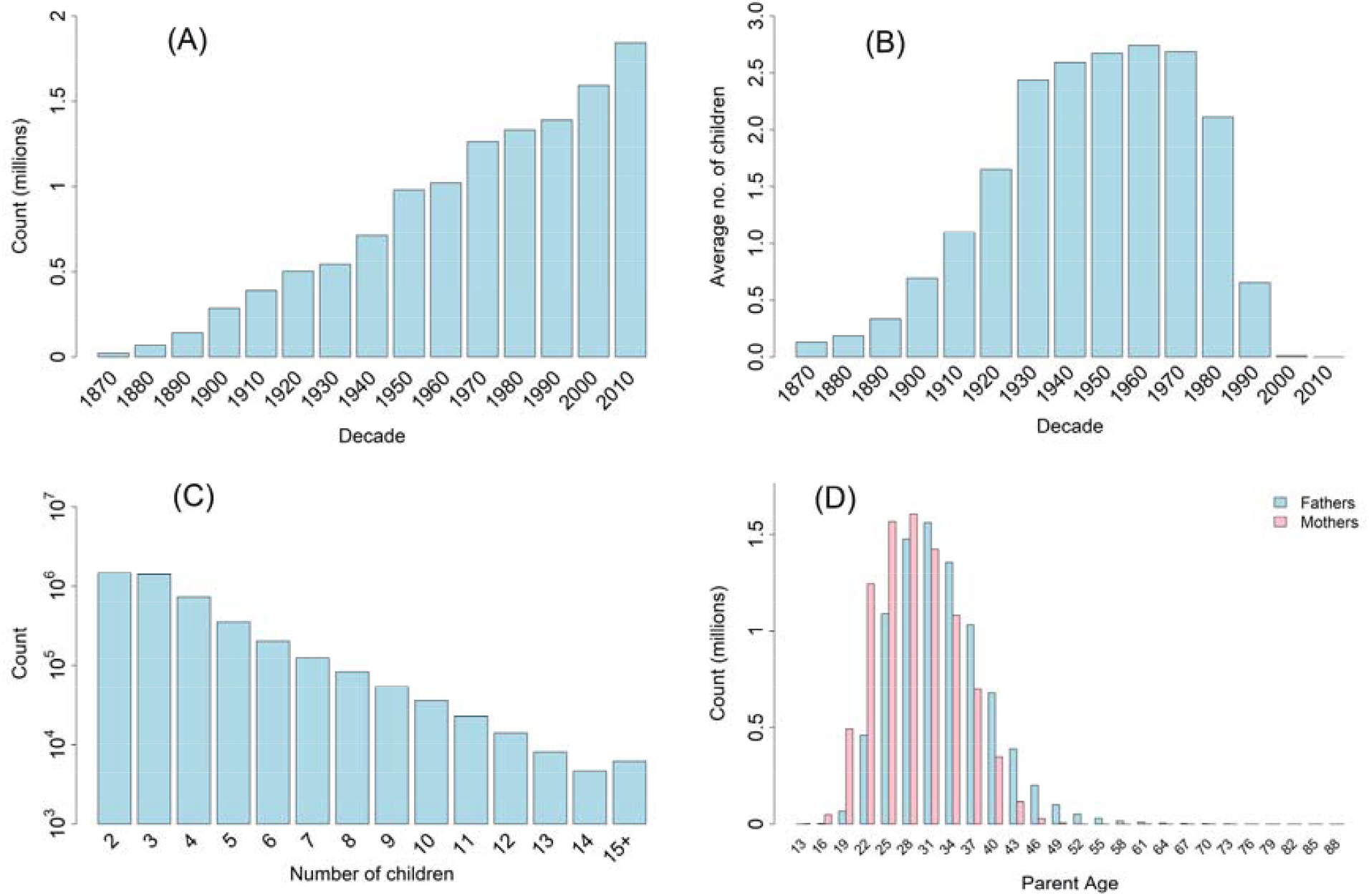
Characteristics of the IPIA data. (A) The distribution of birth decades (1870s-2010s). (B) The average number of children per individual (including childless individuals or individuals who have not yet completed their families) vs the parental birth decade. Birth decades start at the indicated year (e.g., the 1960 birth decade corresponds to birth years 1960-1969). (C) The distribution of the number of children per individual. We combined all individuals with 15 or more children into a single category (see also Figure S1). The y-axis is logarithmic. (D) The distribution of the paternal (light blue) and maternal age (pink) at childbirth, also known as the generation interval. The distribution is over all births in the database. Numbers on the x-axis indicate the midpoints of 3-year bins (e.g., 25 corresponds to ages 24-26).

To simulate FIGG, we designated a random subset of the living individuals as the “genomic database”, and recorded the fraction of the population with at least one genetic relative in the database (see Methods). We call this fraction the “match rate”. While we cannot identify actual genetic relatives (given the lack of genetic data), we assume that all relationships up to 7^th^-degree (third-cousins or equivalent; see Table S2 for definitions and examples) could be reliably detected^19–22^. We note that even given a documented path in the population register connecting the target person to a relative in the genomic database, identification is not guaranteed. Therefore, our estimated match rate is not equal to the identification rate (see the Discussion). We repeat the simulation several times for each database size, showing the average match rate vs the database size for a number of values for the maximal permitted relationship (Figure 2).

**Figure 2.**
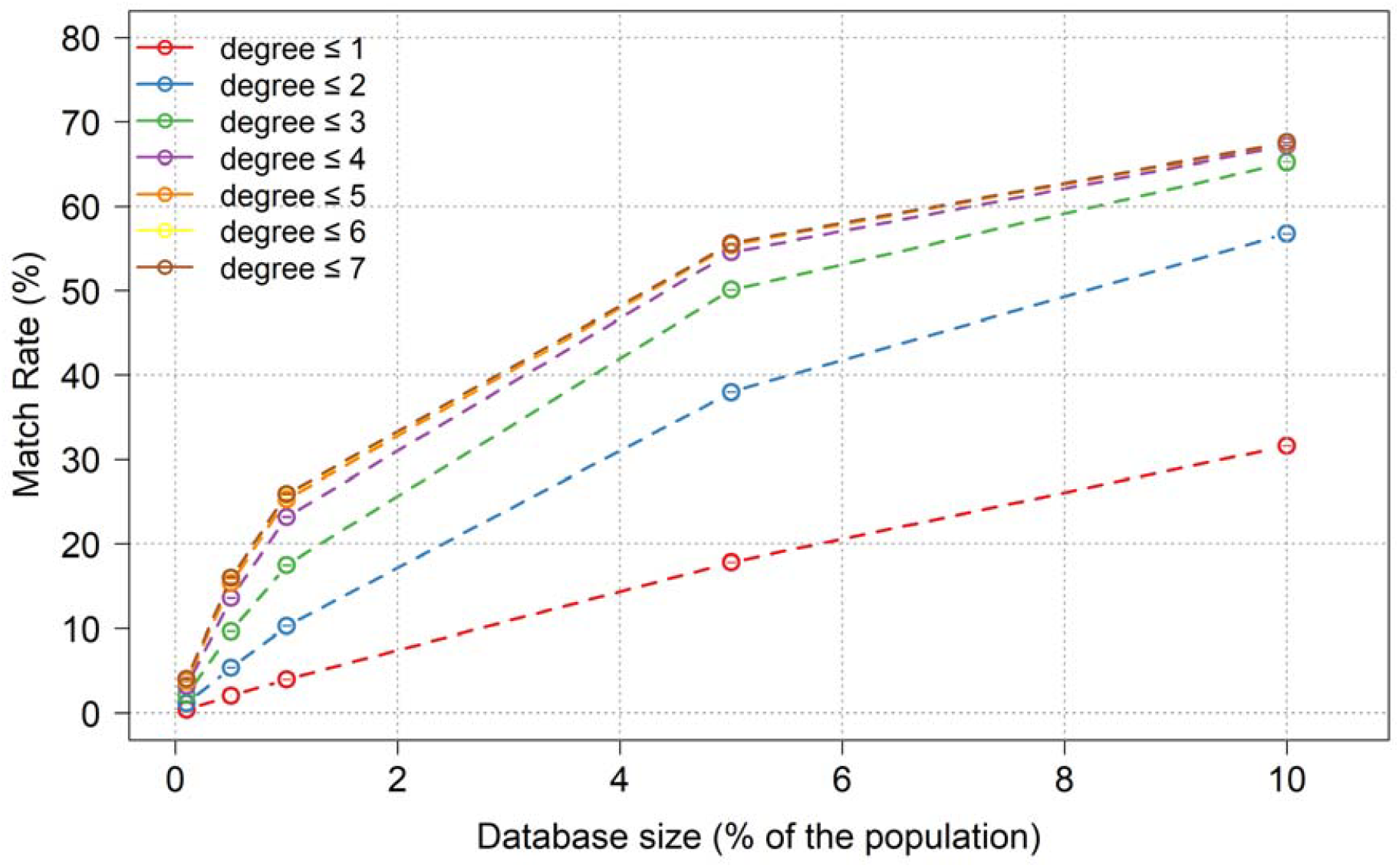
The FIGG match rate using a national population register. We plot the proportion of the Israeli population (including deceased) who have at least one relative in the genomic database to whom they connect via a path in the population register of a maximal pre-defined length. The x-axis is the database size, in proportion of the living population over 18 years old. Each curve (legend) corresponds to a different maximal degree of relationship. For example, a first-degree relative is a parent or a full sibling, and a second-degree relative can be a half-sibling, a grandparent, or an aunt. Each data point is the average over 100 simulations. Standard deviations across simulations are shown as error bars, but are too small to be visible. All data used to generate the figure appears in Data S1.

The results show that with a database covering 1% of the population, the match rate, considering relationships up to 7^th^-degree, is around 26%. Databases covering 5% or 10% of the population reach a match rate of 56% and 68%, respectively. Thus, once databases accumulate a critical mass of genomes, a considerable fraction of the population is potentially amenable to identification. The results are similar across a number of sensitivity analyses: removing extremely fertile individuals (≥15 children); including deceased individuals in the database; and limiting the targets only to those alive (Figure S2).

Surprisingly, the match rate in Figure 2 barely increases between 5^th^-degree (e.g., second cousins) and 7^th^-degree (e.g., third cousins) relatives. For example, for a database covering 1% of the population, the match rate increases from 25.3% with 5^th^-degree relatives to just 25.9% with 7^th^-degree relatives. This result, which is unexpected based on theoretical expectations^2,12^, is likely due to the limited timespan of the register; given that Israel was only founded in 1948, very few pairs of individuals have a common great-great-grandparent in the register. Consequently, the vast majority of individuals with a match in the database have a close relative there. For example, databases covering 5% or 10% of the population guarantee that 50% or 65% of targets, respectively, have a *3*^*rd*^-*degree* (e.g., a first-cousin or a great-grandparent) or a closer relative.

FIGG cases can be difficult to solve with just a single database match, given the very large number of potential relatives of the match that must be examined^2^. In practice, many cases are solved by identifying a common ancestral couple for each cluster of matches and determining how the family trees for the clusters intersect^4,17^. In Figure 3A, we show the match rate when requiring at least two genetic relatives in the database. With a database covering 1% of the population, the “double” match rate (5^th^-degree or closer relatives) is 9.4%, while it is 37% and 51% for databases covering 5% and 10% of the population, respectively. Focusing on a database covering 1% of the population, we show in Figure 3B the distribution of the number of database matches among target individuals with at least one match (7^th^-degree or closer). We find that 37% of those targets have more than one match, 15.5% have three or more matches, and 7.1% have four or more. Thus, requiring multiple matches with a genealogical relationship documented in the register reduces the expected FIGG success rate or requires larger databases.

**Figure 3.**
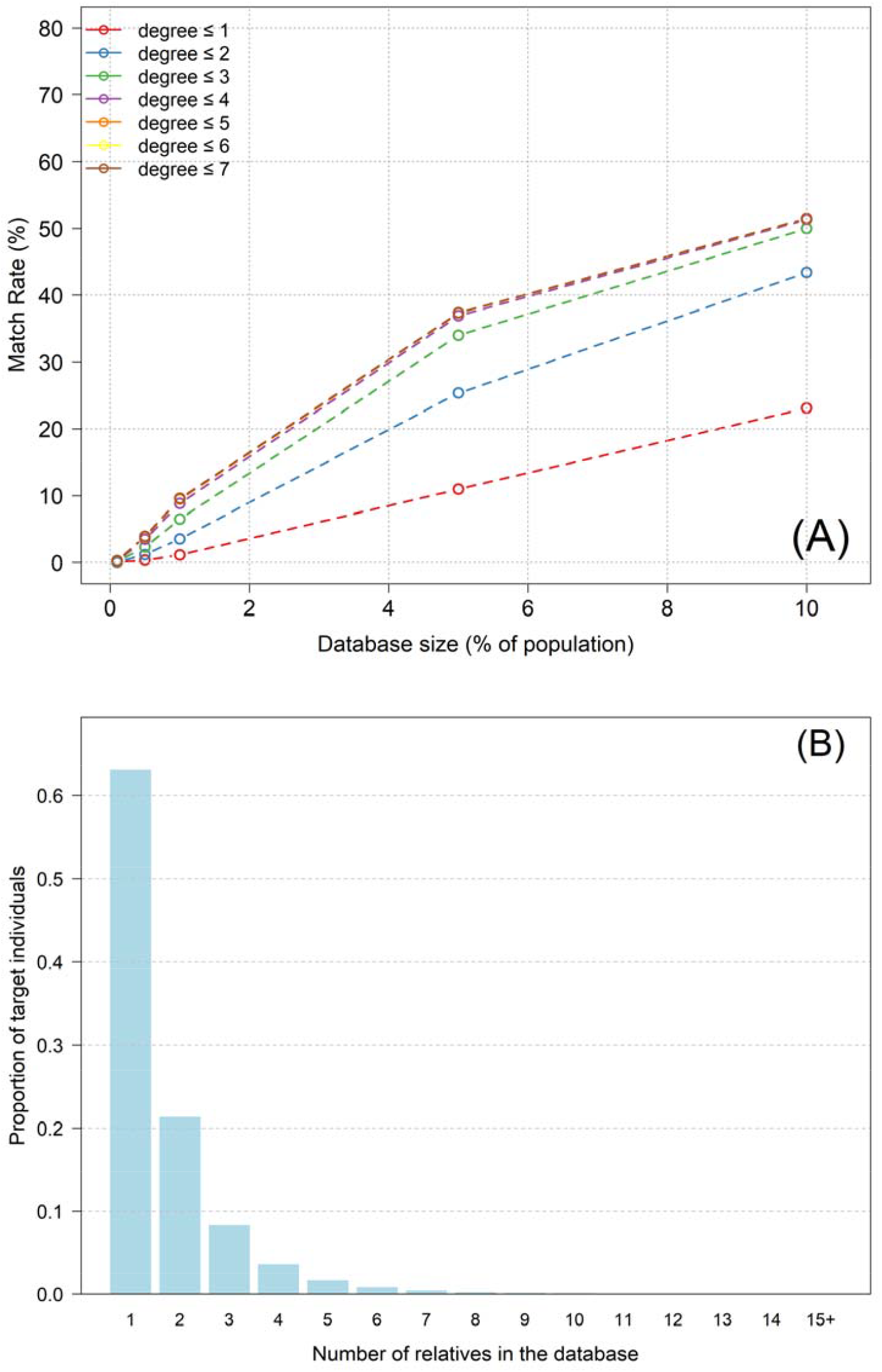
The probability of detecting multiple relatives of the target in the database. (A) The match rate when requiring at least two relatives in the database. All parameters are as in Figure 2. (B) The distribution of the number of relatives in the database. We only considered target individuals (alive or deceased) who have at least one relative in the database. The database size was 1% of the population (alive and >18) and we considered all 7^th^-degree or closer relatives. Target individuals *already* in the database count themselves towards the total number of relatives that they have there.

## Discussion

In this work, we leveraged a unique national resource to realistically simulate some of the elements of a FIGG search. Specifically, using the Israeli population register, we estimated the proportion of target individuals with genealogically documented relatives (“matches”) in a genomic database. The strength of our data is that it represents the type of genealogical information that is readily available to law enforcement in some countries^18,23,24^. Thus, our estimates reflect “high quality” matches, in the sense that the target person is immediately visible to law enforcement once the match has been detected. This is particularly important given recently imposed restrictions on FIGG by commercial genealogy services^15^ and given the availability of national registers in several countries^25–27^. In countries that permit the use of population registers for FIGG, law enforcement may even search the register as a first-line approach.

Observing a match in a real-life FIGG case does not guarantee successful identification. First, reviewing all relevant relatives of a single match can be extremely tedious, even with a population register. Second, many relatives are likely to be missed from either national registers or public genealogical records. Third, even with multiple matches, their family trees may be impossible to fully reconstruct, leading to no intersection. Targets with multiple matches documented in the register should be relatively easy to identify; however, the number of such targets is relatively small (Figure 3). Finally, unless a descendant relative is found, siblings cannot be distinguished using the FIGG approach without additional forensic information.

Our approach for estimating the match rate has several additional limitations. First, clerical errors are inevitable in a database developed over more than seven decades. Second, some recorded child-parent relationships may not be genetic, due to non-paternity, adoption, gamete donation, hospital mix ups, etc. Third, the limited time depth of the Israeli register severely limits the number of documented distant relatives, underestimating the potential real-life match rate. Given the occasional availability of external genealogical records that reach deeper into the past compared to the register, we expect the empirical FIGG match rate to be somewhat higher than estimated here. Fourth, some genetic relatives will be missed by current genomic methods, and some non-relatives will be wrongly identified as relatives. However, this is expected to be rare for third cousins and extremely rare for second cousins or closer. Fifth, more commonly, a relationship may be inferred, based on the genetic data, to be of a different degree or a different type (e.g., a first cousin instead of a great uncle) compared to the genealogical relationship. Families with monozygotic twins would also show discordance between genetic and genealogical relationships. In endogamous and consanguineous populations, individuals are often related via multiple paths, making the genealogical relationship less well defined. However, the effect on our match rate estimates is expected to be small, because relatives should be genetically detectable regardless of this limitation. Sixth, it is assumed that the quality of the DNA from the forensic sample is sufficiently high, such that all genetic matches can be trusted. Finally, it is unclear how the results will generalize to other countries, which may differ in their fertility rates over time, their mating and consanguinity patterns, and the depth of their genealogical records.

Our study of the Israeli population establishes the expected FIGG match rate for genealogically documented relatedness across different database sizes, genetic relationships, and minimal numbers of matches. This information is important for policymakers regulating databases or FIGG, as well as to those planning to build databases for FIGG or use FIGG in specific cases. The results also underscore privacy risks in countries where large-scale genomic databases are available. Existing and emerging biobanks in multiple countries, including the UK^28,29^, Finland^30^, Iceland^31^, Estonia^32^, Taiwan^33^, Singapore^34^, the UAE^35^, and Qatar^36^, already comprise large proportions of their respective populations. Further, direct-to-consumer genomic databases include tens of millions of individuals and are vulnerable to market upheavals^37^. Our results quantify the privacy risks of unauthorized access to these large-scale databases.

## Methods

### Ethics and approvals

The study was approved by the research committee of the Israel Central Bureau of Statistics (ICBS), study no. 16810376. The data was provided to the ICBS by the IPIA. The IPIA also approved the study. The data was de-identified by replacing each ID with a random number. No statistics were reported for any category with less than ten individuals. All analyses were performed on the ICBS research platform on computers disconnected from the internet and using files prepared specifically for this project. All outputs were reviewed by ICBS staff.

### The genealogical data

The IPIA dataset includes one row for every past and present citizen of Israel. For each individual, the dataset includes self ID, father’s ID, and mother’s ID. Whenever one or both parents of an individual were not present in the data, no ID was provided. For de-identification, each ID was converted to the same random number across all instances of that ID. Additional information per individual included year of birth, sex (M/F/Unknown), and whether alive (Y/N). The data was up-to-date as of the end of 2021. The dataset includes Israeli citizens living abroad, which is why the number of living people in the dataset is larger than the current Israeli population size.

### Quality control and basic statistics

We generated the list of children of each individual by concatenating the IDs of all individuals who had the focal person as their parent. We then performed the following quality control analyses. In all filters described below, the number of individuals removed is with respect to the original dataset, and some individuals have been removed due to multiple reasons.

#### Illogical or missing data

We verified that all rows had a unique individual ID and that no individual had parents with the same ID^26^. We identified and removed 51,368 individuals with an unknown sex, 3,368 females who were listed as fathers, and 1,289 males who were listed as mothers. We also removed 2,245 individuals listed as both fathers and mothers. Next, we removed 189 individuals who were part of a pedigree “loop”, where an individual is recorded as its own ancestor^38^. We also removed 3,523 individuals without year of birth (among whom only ≤10 had their parents listed). There were 62,346 individuals with a parent whose ID did not have its own row. These parents likely correspond to Palestinian citizens, per IPIA policy, and these records were not removed.

#### Year of birth

We removed 213 individuals who were born between 1800-1849, who would have been over 98 years old at the foundation of Israel in 1948. We also removed 40,239 individuals who were recorded as alive (as of 2021) and were born before 1907. There were 138,918 pairs of full-siblings, 4,106 triplets and 193 groups of four or more siblings born in the same year. There were 7,873 families in which half-siblings with a shared *father* were born in the same year, among them 7,717 involved two mothers and 156 three or four mothers. In contrast, there were 238 families in which half-siblings with a shared *mother* were born in the same year. None of the above was removed. We only excluded ≤10 mothers who, implausibly, had children born in the same year with three different fathers.

#### Number of children

There were 92,364 individuals with 10 or more children (54.3% male) and 6,248 with 15 or more children (74.3% male). The distribution of birth decades and the number of children for individuals with 10 or more children is shown in Figure S1. In the absence of additional information, we did not remove any records based on the number of children. However, as part of the sensitivity analyses, we excluded the individuals with 15 or more children from some of the FIGG simulations (Figure S2). The proportion of individuals without children in each birth decade is shown in Figure S1, demonstrating that it is lowest for people born between 1930-1980 (under 40%). There were 6,765 men who had children with three or more women, and 1,760 women who had children with three or more men. We did not remove these individuals.

#### Generation interval

The age at child birth was computed as the difference between the integer years of birth of the parent and the child. The distribution of the paternal and maternal ages at birth is shown in Figure 1, and the mean age at birth vs the birth decade is shown in Figure S1. We removed 81 individuals whose fathers were over 90 years old at birth and 5058 individuals whose mothers were over 50 years old at birth (given that in most such cases, even if real, the mother is most likely not genetically related to the child). We removed 5041 individuals whose parents, at the time when they gave birth, were younger than 12 years old (or were not yet born based on the records).

#### Parental relatedness

We removed 1361 pairs of parents who were siblings, among them 81 were full siblings.

Overall, we removed 105,895 individuals from the IPIA dataset, leaving 12,457,725 individuals for the downstream analyses.

### Simulating FIGG

To simulate FIGG, we first selected a random subset of individuals to form the “genomic database”. In the main analysis, these individuals were required to be alive and over 18 years old (born in or before 2003). We also performed a sensitivity analysis in which the “database individuals” were selected entirely at random, to represent a decades-long process of genomic database development (Figure S2). We assumed that law enforcement investigators have access to genome-wide data for all database individuals. For each genomic database size (0.1%, 0.5%, 1%, 5%, and 10%), we repeated the simulation 100 times, each time selecting a different random group of database individuals.

In a hypothetical FIGG setting, we have the DNA profile of a target individual, whom we attempt to identify by finding genetic relatives (“matches”) in the genomic database (Figure S3). We define the match rate as the proportion of targets with at least one relative in the database. We only consider relationships up to *d*_max_ = 7^th^ degree, such as third cousins, as more distant relatives often share no DNA, while third cousins or closer relatives are very rarely missed^19–22^. The degree of relationship *d* corresponds to a kinship coefficient 2^-*d*^ /2 (see Table S2 for examples). To find the number of people with at least one match in the database, we go over each database individual in turn, and add all of their relatives to the list of targets with matches. Note that the database individuals themselves also have a match, by virtue of having a “zero degree” relative (=self) in the database. In the main analysis, all individuals, including deceased, were possible targets. In a sensitivity analysis, we only attempted to identify living people (Figure S2).

For each database individual, a path to a relative must have zero or more steps “up” (to a parent), then zero or one “side” steps (to a sibling), and finally zero or more steps “down” (to a child). A step towards a half-sibling increases the degree of relationship by 2. See Figure S4 for a visualization.

Accordingly, to create a list of all relatives of a focal person, we first add all ancestors of that person (“up” steps), up to a maximum of *d*_max_ generations back. For each ancestor reached, we record the number of steps that was required to reach it, and then add the set of all of their siblings (“side” step). Then, for each person in the list, we add all of their descendants (“down steps”), as long as the total number of steps from the focal individual is *d*_max_ or less. For each relative, we ignore all relationships with the focal individual except the closest. After performing the search starting from each of the database individuals, we record the number of uniquely reachable relatives.

For the analyses of Figure 2, if an individual is related to more than one database individual, we only retain the closest relationship. To plot the match rates in Figure 2, we average the proportion of targets with at least one relative in the database over all simulations. We also computed the standard deviation of the match rate across simulations and plotted them as error bars (which are not visible given how small they are; see data in Data S1). Similar simulations and plots were generated for the sensitivity analyses. For the analyses of Figure 3, we also record, for each target person, to how many database individuals it is related.

## Acknowledgements

We thank Amy Williams for discussions and Yifat Klopstock from the ICBS for assistance with generating the dataset and reviewing the outputs.

## Data availability

Data used for this study was provided by the ICBS. Other researchers can submit a data access request at https://www.cbs.gov.il/he/CBSNewBrand/Pages/%D7%97%D7%93%D7%A8-%D7%9E%D7%97%D7%A7%D7%A8.aspx.

## Code availability

Code used to generate all the results reported in this study is available at https://github.com/harelg10/forensic-genetic-genealogy-simulation.

## Author contributions

Conceptualization: SC, GG; Formal Analysis: HI, SC; Funding Acquisition: SC; Investigation: HI, SC; Methodology: SC; Project Administration: SC; Software: HI; Supervision: SC; Visualization: HI, SC; Writing – Original Draft Preparation: SC; Writing – Review & Editing: SC, GG, DK, HI.

## Supplementary Figures

**Figure S1.**
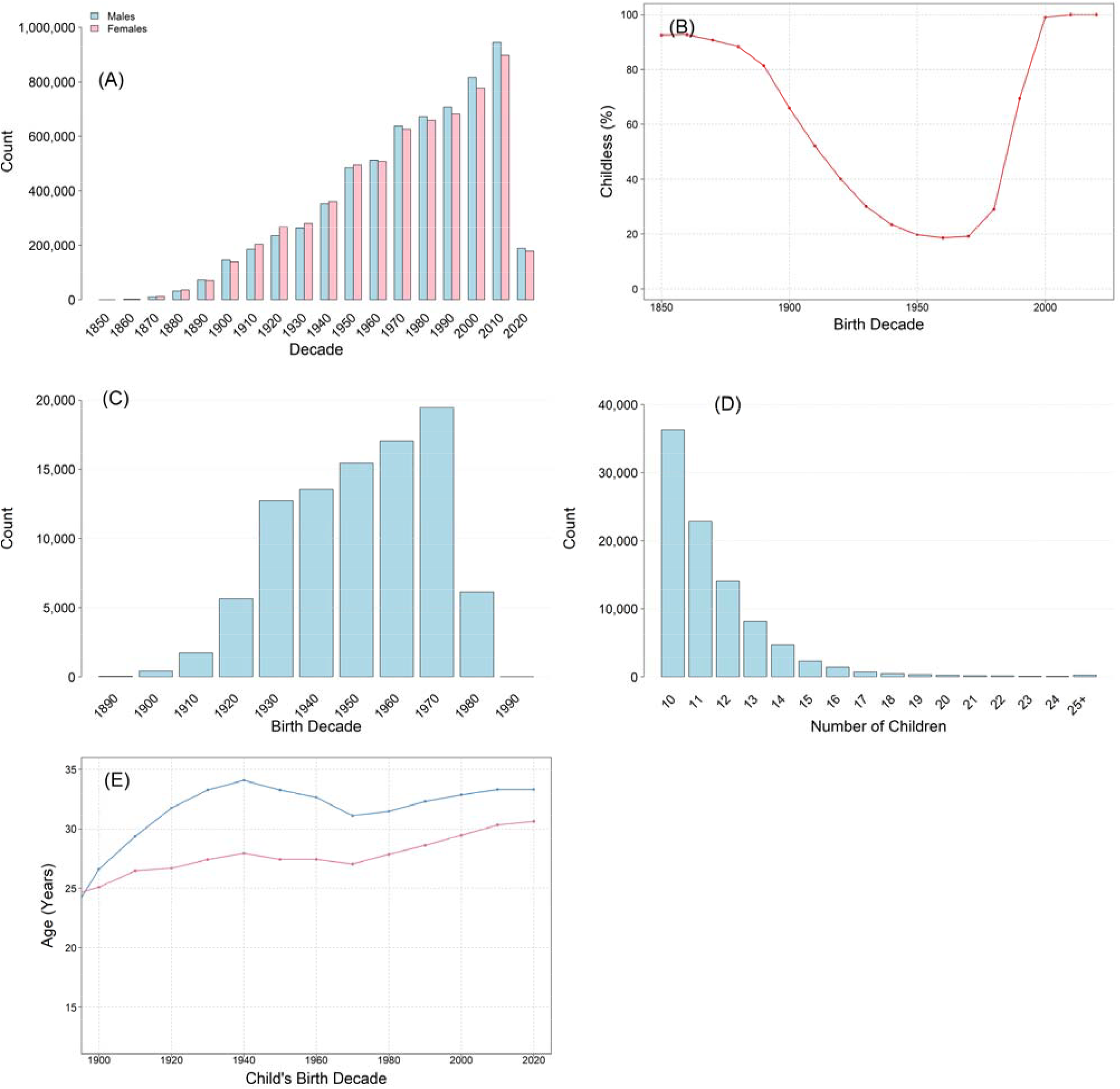
Additional characteristics of the IPIA data. (A) The distribution of birth decades per sex. (B) The proportion of individuals in each birth decade without children. (C) The distribution of the birth decade for individuals with 10 or more children. (D) The distribution of the number of children for those with 10 or more children. All individuals with 25 or more children were grouped into a single bin. (E) The mean paternal and maternal age at birth across birth decades. Each point is an average over all recorded births in the database from the given decade. Birth decades start at the indicated year (e.g., the 1960 birth decade corresponds to birth years 1960-1969).

**Figure S2.**
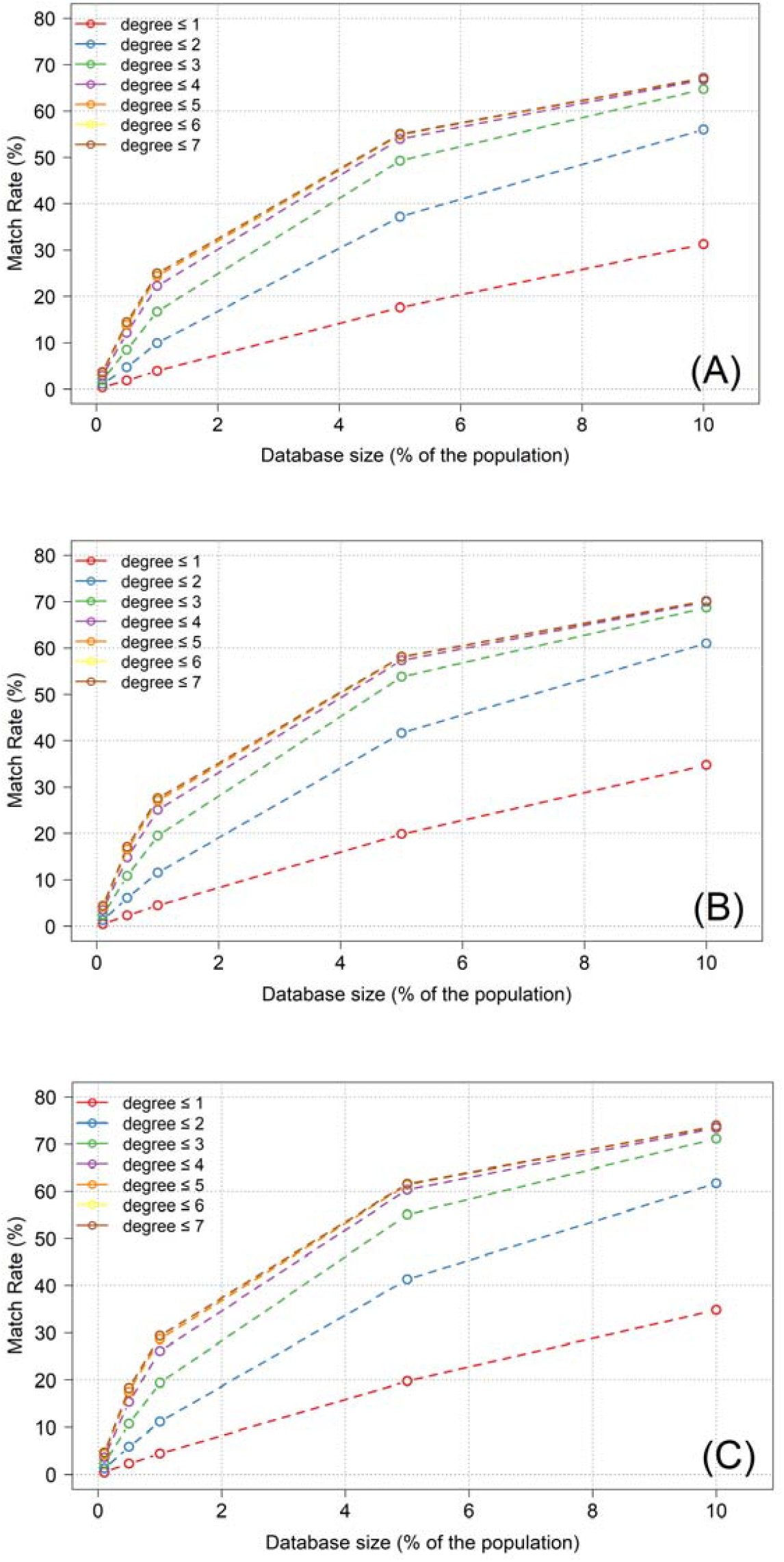
Sensitivity analyses for the estimated FIGG match rates. All parameters were as in the main analysis (Figure 2), except the following. (A) Removing individuals with 15 or more children. (B) Assuming that already deceased individuals can also be selected to the database. (C) Assuming that only living individuals are targets. In each panel, we plot the average proportion of the targets with at least one relative in the database, for a number of values for the maximal degree of relationship, as in Figure 2.

**Figure S3.**
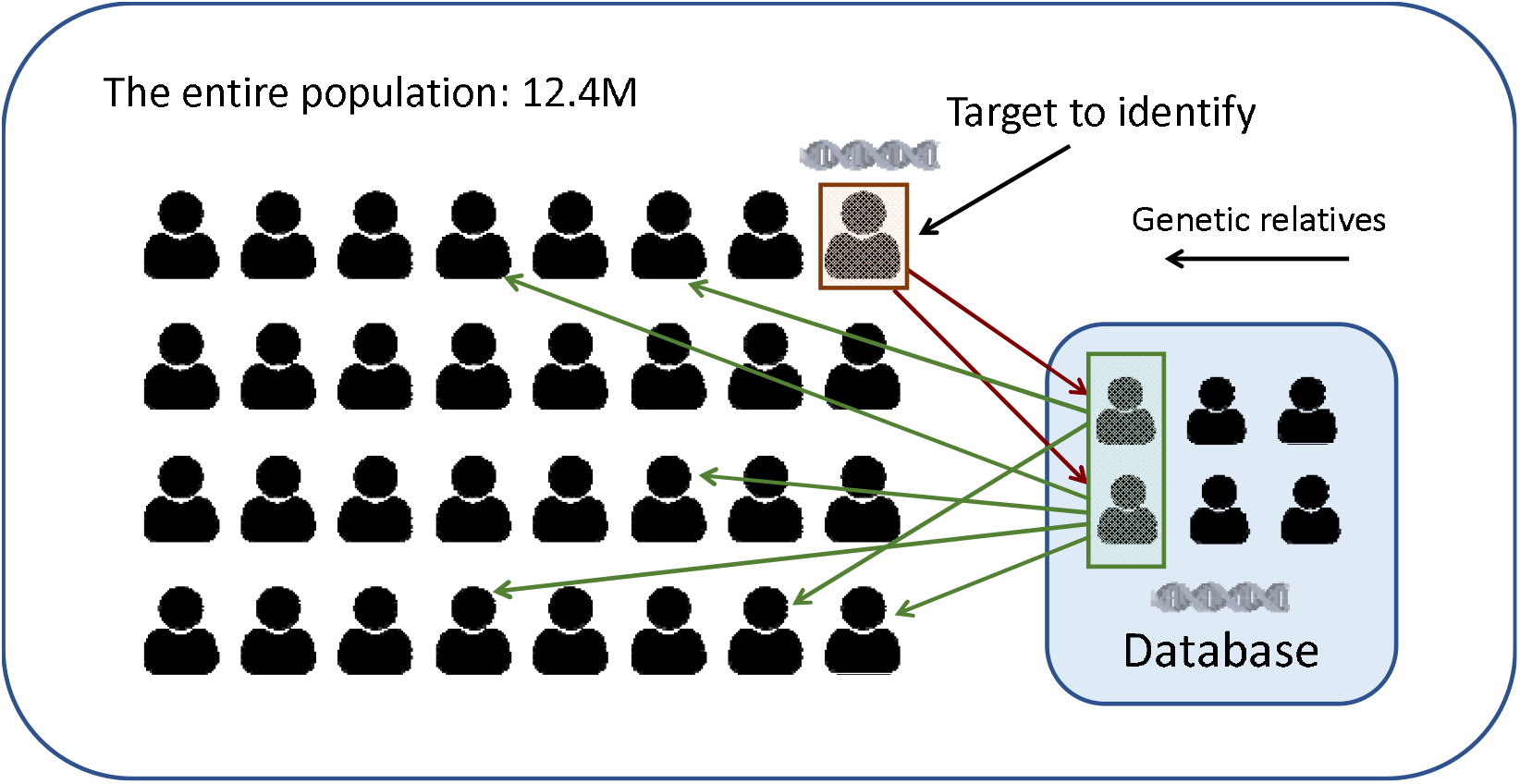
A schematic of FIGG. For a subset of the population, a genome-wide DNA profile is available in a “genomic database” (light blue box). The identity of a target person (peach box) is unknown, but we have access to their genome-wide genetic profile. By comparing the DNA profile of the target against that of all individuals in the database, genetic relatives of the target (“matches”; red arrows) can be identified (light green box inside the database). Once matches have been detected, the investigation focuses on their relatives (green and red arrows), given that the target person must be one of them. Thus, to be identifiable using FIGG, a target person must have at least one relative in the database. We require the relationship to be equal or closer to a prespecified degree, corresponding to the most distant relationship that can be confidently detected using genetic data (usually 7^th^ degree, e.g., a third cousin). In practice, more distant relationships can also sometimes be detected.

**Figure S4.**
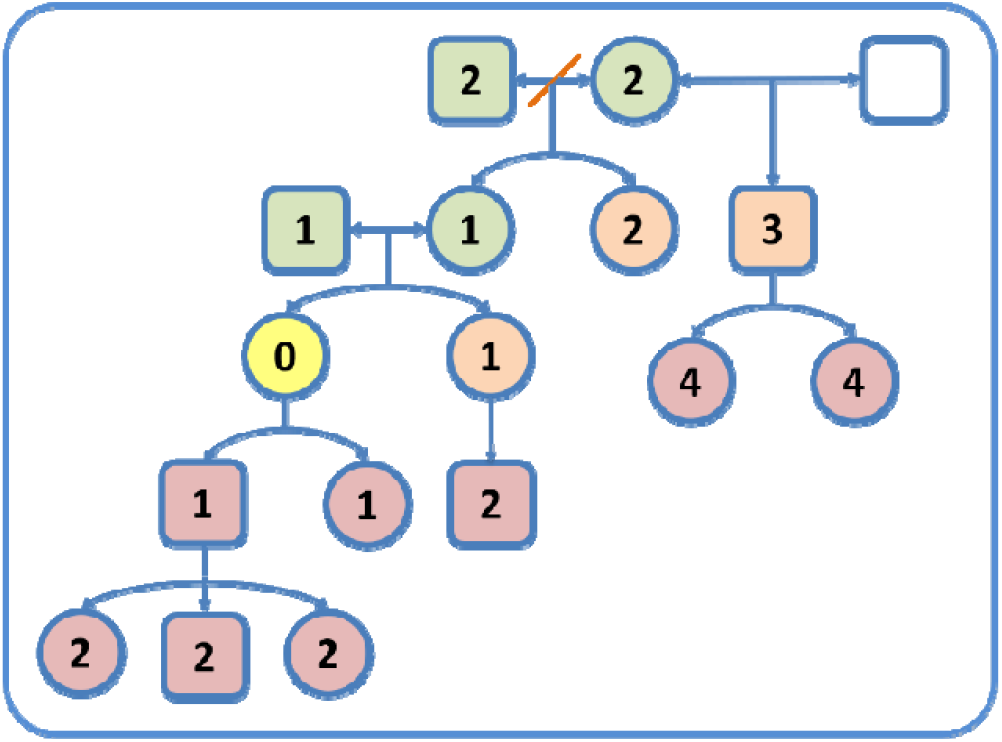
A visualization of the search for all relatives of a focal individual. In the pedigree diagram, squares are males and circles are females. The number inside each shape indicates the degree of relationship to the focal individual. Accordingly, the focal individual is marked as having degree 0 and is colored yellow. The other colors represent the type of “step” that was taken to reach the individual: steps “up” are colored light green; “side” steps peach, and “down” steps pink. The red diagonal line indicates a “divorce”. The child from the second marriage is a half-sibling of the focal individual’s mother, and its degree of relationship to the focal individual is thus 3. Not all members of the pedigree are shown.

## Supplementary Tables

**Table S1.** Statistics of the IPIA dataset before and after QC. Rows are the same as in Table 1.

| Statistic | Before QC | After QC |
| --- | --- | --- |
| Total number of individuals | 12,563,620 | 12,457,725 |
| Number of individuals alive | 10,498,709 | 10,430,877 |
| Number of deceased individuals | 2,064,911 | 2,026,848 |
| Number of males | 6,288,657 | 6,262,793 |
| Number of females | 6,223,595 | 6,194,932 |
| Number of individuals with each number of parents recorded | 0 parents: 3,553,767<br>1 parent: 692,430<br>2 parents: 8,317,423 | 0 parents: 3,465,674<br>1 parent: 689,477<br>2 parents: 8,302,574 |
| Number of individuals with one or more half-siblings | 690,460 | 685,856 |
| Fraction of individuals without children | 55.52% | 55.24% |
| Mean (SD) of the number of children per individual | All individuals: 1.37 (2.04)<br>Only individuals with children: 3.08 (2.01) | All individuals: 1.38 (2.04)<br>Only individuals with children: 3.09 (2.01) |
| Mean (SD) of the paternal age at birth | 32.5 (6.85) | 32.5 (6.78) |
| Mean (SD) of the maternal age at birth | 28.7 (5.89) | 28.7 (5.82) |

**Table S2.** Examples for relationships of different degrees. The degree of relatedness d corresponds to a kinship coefficient 2^-d^/2. The coefficient is defined as the probability of two random alleles, one from each individual, to be identical by descent (i.e., derive from the same ancestor). The kinship coefficient is itself half of the proportion of the genome shared by descent. The examples assume no consanguinity (relatedness between the parents), and that the individuals are not related via additional paths. Note that the number of meioses separating the individuals does not define the degree of relatedness. For example, a parent and a child are separated by a single meiosis (the parent-to-child transmission), while full-siblings are separated by two meioses (parent to both children); nevertheless, both pairs are first-degree relatives.

| Degree of relationship | Kinship coefficient | Examples |
| --- | --- | --- |
| 0 | $2^{-0}/2 = 1/2$ | Self, identical twin |
| 1 | $2^{-1}/2 = 1/4$ | Parent, child, full-sibling |
| 2 | $2^{-2}/2 = 1/8$ | Grandparent, grandchild, half-sibling, uncle/aunt, nephew/niece, double first cousins |
| 3 | $2^{-3}/2 = 1/16$ | Great-grandparent, great-grandchild, granduncle/aunt/nephew/niece, first cousin, half-uncle/aunt/nephew/niece |
| 4 | $2^{-4}/2 = 1/32$ | First cousin once removed, half first cousin, great-great-grandparent/child |
| 5 | $2^{-5}/2 = 1/64$ | Second cousin, first cousin twice removed |
| 6 | $2^{-6}/2 = 1/128$ | Second cousin once removed, half second cousin |
| 7 | $2^{-7}/2 = 1/256$ | Third cousin, second cousin twice removed |

